# Covert spatial attention is uniform across cardinal meridians despite differential adaptation

**DOI:** 10.1101/2025.09.30.676904

**Authors:** Hsing-Hao Lee, Marisa Carrasco

## Abstract

Visual adaptation and attention are two processes that help manage the brain’s limited bioenergetic resources for perception. Visual perception is heterogeneous around the visual field: it is better along the horizontal than the vertical meridian (horizontal-vertical anisotropy, HVA), and better along the lower than the upper vertical meridian (vertical meridian asymmetry, VMA). Recently, we showed that visual adaptation is more pronounced at the horizontal than the vertical meridian, but whether and how this differential adaptation modulates the effects of covert spatial attention remains unknown. In this study, we investigated whether and how the effects of endogenous (voluntary) and exogenous (involuntary) covert attention on an orientation discrimination task vary at the cardinal meridians, with and without adaptation. We manipulated endogenous (Experiment 1) or exogenous (Experiment 2) attention via an informative central or uninformative peripheral cue, respectively. Results showed that (1) in the non-adapted condition, the typical HVA and VMA emerged in contrast thresholds; (2) the adaptation effect was stronger at the horizontal than the vertical meridian; and (3) regardless of adaptation, both endogenous and exogenous attention enhanced and impaired performance at the attended and unattended locations, respectively, to a similar degree at both cardinal meridians. Together, these findings reveal that, despite differences between endogenous and exogenous attention, their effects remain uniform across cardinal meridians–even under differential adaptation that reduces intrinsic asymmetries of visual field representations.

## Introduction

Visual adaptation and attention are two processes that optimize performance and help manage the brain’s limited bioenergetic resources by allocating them according to task demands (Carrasco, 2011; Lee, Fernández, & Carrasco, 2024; Lennie, 2003; Pestilli, Viera, & Carrasco, 2007). Although both processes modulate sensory responses, they have opposite effects on the contrast response function. Visual adaptation helps manage bioenergetic resources by increasing metabolic efficiency–it reduces sensitivity to repeated features and enhances sensitivity to novel ones. For example, contrast adaptation can adjust the gain of the neural response so that its dynamic range is matched to the range of levels in the stimulus (Boynton & Finney, 2003; Gardner et al., 2005; Kohn, 2007; Perini, Cattaneo, Carrasco, & Schwarzbach, 2012; Vergeer, Mesik, Baek, Wilmerding, & Engel, 2018; Webster, 2011, 2015). In contrast, visual attention selectively improves information processing at an attended location while impairing processing elsewhere – a ubiquitous performance tradeoff considered a push-pull mechanism (e.g., Dosher & Lu, 2000b; Ling & Carrasco, 2006a; Pestilli & Carrasco, 2005; Pestilli, Ling, & Carrasco, 2009; Pestilli et al., 2007; for reviews, Carrasco, 2011, 2014; Desimone & Duncan, 1995; Olivers, 2025).

There are two types of covert spatial attention: endogenous and exogenous. Endogenous attention is voluntary, goal-driven, and flexible; exogenous attention is involuntary, stimulus-driven, and automatic. Endogenous attention takes ∼300 ms to be deployed and can be sustained for many seconds, whereas exogenous attention peaks at ∼120 ms and is transient (reviews: Carrasco, 2011, 2014). Despite these differences, both types of attention improve performance in many visual tasks, e.g., contrast sensitivity (e.g., Herrmann, Montaser-Kouhsari, Carrasco, & Heeger, 2010; Pestilli et al., 2009), appearance (review: Carrasco & Barbot, 2019), and orientation discrimination (e.g., Fernández, Okun, & Carrasco, 2022). However, they have distinct effects in other tasks, e.g., texture segregation (e.g., Barbot & Carrasco, 2017; Jigo, Heeger, & Carrasco, 2021; Yeshurun & Carrasco, 1998), and alter sensitivity across a different spatial frequency range (Fernández et al., 2022; Jigo & Carrasco, 2020).

Exogenous attention restores contrast sensitivity after adaptation; although adaptation reduces sensitivity, the magnitude of the exogenous attentional benefit at the attended location and its concurrent cost at the unattended location remain comparable to those observed without adaptation (Lee et al., 2024; Pestilli et al., 2007). However, whether and how endogenous attention operates after adaptation is unknown. Thus, our first goal was to examine whether endogenous attention restores contrast sensitivity after adaptation. It is possible that after adaptation endogenous attention (1) enhances contrast sensitivity to a similar extent as without adaptation, assuming that similar to exogenous attention (Lee et al., 2024; Pestilli et al., 2007), endogenous attention and adaptation yield independent effects on contrast sensitivity (**Figure 1A**; Hypothesis 1); (2) enhances sensitivity more than before adaptation, reflecting a compensatory process given the flexible nature of endogenous attention, which optimizes performance as a function of task demands (Barbot & Carrasco, 2017; Barbot, Landy, & Carrasco, 2012; Giordano, McElree, & Carrasco, 2009; Hein, Rolke, & Ulrich, 2006; Yeshurun, Montagna, & Carrasco, 2008); it may help more than without adaptation, as after decreases there is more room for improvement (**Figure 1B**; Hypothesis 2); or (3) enhances sensitivity less than without adaptation; if reduced baseline sensitivity limits the push-pull effects of endogenous attention (**Figure 1C**; Hypothesis 3).

**Figure 1.**
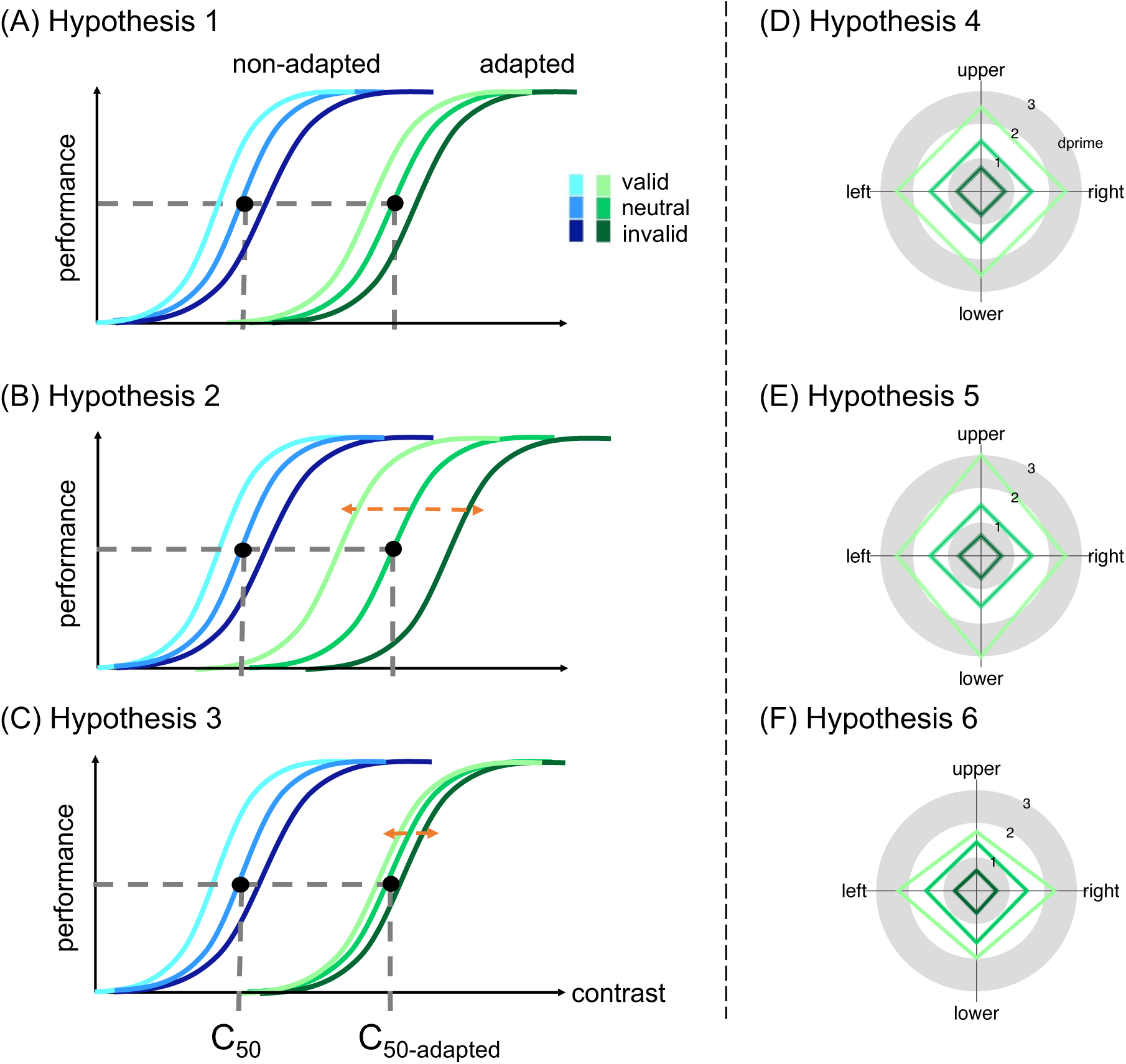
(A-C) Hypotheses regarding effects on contrast sensitivity. (A) Hypothesis 1: Attentional effect is comparable with and without adaptation. The c50 and c50-adapted indicate the contrast threshold derived from the titration procedures in the non-adapted and adapted conditions, respectively. (B) Hypothesis 2: Attentional effect is larger with than without adaptation. (C) Hypothesis 3: Attentional effect is smaller than without adaptation. (D-F) Hypotheses regarding effects on contrast sensitivity as a function of location at the cardinal locations. (D) Hypothesis 4: Attentional effect is comparable around polar angle after adaptation. (E) Hypothesis 5: Attentional effect is stronger at the vertical than horizontal meridian. (F) Hypothesis 6: Attentional effect is smaller at the vertical than horizontal meridian.

Because endogenous attention is flexible (Barbot & Carrasco, 2017; Barbot et al., 2012; Giordano et al., 2009; Hein et al., 2006; Yeshurun et al., 2008), but exogenous attention is not (Barbot et al., 2012; Carrasco, Loula, & Ho, 2006; Crotty, Massa, Tellez, White, & Grubb, 2025; Giordano et al., 2009; Hein et al., 2006; Luck & Thomas, 1999; Yantis & Jonides, 1996; Yeshurun & Carrasco, 1998), and because distinct brain regions are critical for their effect –right frontal eye fields for endogenous attention (Fernández, Hanning, & Carrasco, 2023), and early visual cortex for exogenous attention (Fernández & Carrasco, 2020; Lee et al., 2024), where it interacts with adaptation (Lee et al., 2024) –it is possible that they exert different effects on contrast sensitivity after adaptation. Therefore, our second goal was to determine whether endogenous and exogenous attention have similar or different effects on contrast sensitivity following adaptation.

Finally, we investigated whether target location matters. In adult humans, visual performance is better at the horizontal than the vertical meridian (horizontal-vertical anisotropy, HVA), and better at the lower than the upper vertical meridian (vertical meridian asymmetry, VMA). These visual field asymmetries, known as performance fields, are present in many fundamental visual tasks, including contrast sensitivity (Abrams, Nizam, & Carrasco, 2012; Baldwin, Meese, & Baker, 2012; Cameron, Tai, & Carrasco, 2002; Carrasco, Talgar, & Cameron, 2001; Corbett & Carrasco, 2011; Fuller, Rodriguez, & Carrasco, 2008; Himmelberg, Winawer, & Carrasco, 2020; Lee & Carrasco, 2025; Purokayastha, Roberts, & Carrasco, 2021), visual acuity (Kwak, Hanning, & Carrasco, 2023; Montaser-Kouhsari & Carrasco, 2009), spatial resolution (Altpeter, Mackeben, & Trauzettel-Klosinski, 2000; Carrasco, Williams, & Yeshurun, 2002; Greenwood, Szinte, Sayim, & Cavanagh, 2017; Talgar & Carrasco, 2002), and motion (Fuller & Carrasco, 2009; Tünçok, Kiorpes, & Carrasco, 2025), as well as mid-level visual processes, such as texture segregation (Barbot, Xue, & Carrasco, 2021; Greenwood et al., 2017; Talgar & Carrasco, 2002; Z. Wang, Murai, & Whitney, 2020) and crowding (Greenwood et al., 2017; Kurzawski et al., 2023; Petrov & Meleshkevich, 2011), and high-level tasks, such as numerosity perception (Chakravarthi, Papadaki, & Krajnik, 2022), face perception (Afraz, Pashkam, & Cavanagh, 2010; Kim & Chong, 2024), word identification (Tsai, Liao, Hou, Jang, & Chen, 2024), and visual short-term memory (Montaser-Kouhsari & Carrasco, 2009).

These visual field asymmetries are resistant to endogenous (Purokayastha et al., 2021; Tünçok, Carrasco, & Winawer, 2025) and exogenous attention (Cameron et al., 2002; Carrasco et al., 2001; Roberts, Ashinoff, Castellanos, & Carrasco, 2018; Roberts, Cymerman, Smith, Kiorpes, & Carrasco, 2016), as well as to temporal (Fernández, Denison, & Carrasco, 2019) attention. Thus, performance fields are not easily reshaped. On the contrary, presaccadic attention, which enhances the processing at the location of the impending saccade target, exacerbates performance asymmetries at the cardinal locations by enhancing contrast sensitivity the most at the horizontal meridian and the least at the upper vertical meridian (Hanning, Himmelberg, & Carrasco, 2022, 2024; Kwak, Hanning, & Carrasco, in press; Kwak, Zhao, Lu, Hanning, & Carrasco, 2024).

A recent study showed that visual adaptation is stronger at the horizontal than the vertical meridian, leading to more homogeneous perception by mitigating the HVA (Lee & Carrasco, 2025). It remains unknown, however, whether and how endogenous and exogenous attention reshape performance fields after such differential adaptation. Thus, our third goal was to investigate whether, following adaptation, covert spatial attention enhances contrast sensitivity (1) to the same extent at the cardinal meridians around polar angle, similar to without adaptation (e.g., Carrasco et al., 2001; Purokayastha et al., 2021; Roberts et al., 2018; Roberts et al., 2016; Tünçok, Carrasco, & Winawer, 2025) (**Figure 1D**; Hypothesis 4), (2) more at the vertical than the horizontal meridian, and more at the upper than the lower vertical meridian, acting as a compensatory mechanism to reduces asymmetries (**Figure 1E**; Hypothesis 5), or (3) more where baseline performance is already better (i.e., the horizontal meridian) than where it is worse (i.e., vertical meridian, especially the upper vertical meridian), thereby exaggerating asymmetries (**Figure 1F**; Hypothesis 6).

Both adaptation (Altan, Morgan, Dakin, & Schwarzkopf, 2025; Dao, Lu, & Dosher, 2006; Gardner et al., 2005; Perini et al., 2012; Pestilli et al., 2007) and endogenous attention (Dosher & Lu, 2000a; Ling & Carrasco, 2006a; Lu, Lesmes, & Dosher, 2002; Pestilli et al., 2009) primarily affect the contrast gain of the contrast response function (**Figure 1A**), i.e., a shift in threshold, whereas exogenous attention primarily affects response gain (Fernández & Carrasco, 2020; Pestilli et al., 2009), i.e., a shift in asymptote. Additionally, according to a prominent normalization model of attention (Reynolds & Heeger, 2009), exogenous attention can also affect contrast gain when the attentional window is wider than the stimulus size, and endogenous attention can also affect response gain when the attentional window is narrower than the stimulus size (Herrmann et al., 2010). In this study, to directly compare the two types of attention before and after adaptation at the cardinal meridians, we induced a larger attentional window in the exogenous attention experiment, enabling contrast gain effects predicted by Reynolds and Heeger’s (2009) normalization model of attention.

In summary, we asked (1) whether and how endogenous attention restores contrast sensitivity following adaptation, (2) whether endogenous and exogenous attention have similar or distinct effects on contrast sensitivity before and after adaptation, and (3) whether these effects uniformly or differentially across the cardinal meridians around the visual field. These findings are essential for elucidating how the visual system engages adaptation and attention—two fundamental visual processes that manage limited bioenergetic resources—to optimize performance across locations that differ in intrinsic discriminability and in their corresponding representation in cortical surface area.

## Experiment 1 – Endogenous attention

### Methods

#### Participants

Twelve adults (5 females, age range: 24-36 years old), including author HHL, participated in the experiment. All of them had normal or corrected-to-normal vision. Sample size was based on previous studies on adaptation (Lee et al., 2024), with an effect size of *d*=1.3, and on performance fields (Lee & Carrasco, 2025), with an effect size of *d*=1.41 for performance in the neutral trials. According to G*Power 3.0 (Faul, Erdfelder, Lang, & Buchner, 2007), we would need 9 participants for adaptation and 8 participants for performance fields to reach a power=0.9. We also estimated the required sample size for the interaction between adaptation and location, based on a recent study between adaptation and performance fields (Lee & Carrasco, 2025) (*ηp^2^*=0.34), by assuming SD=1, we would need 10 subjects to reach a power=0.9 according to the Monte-Carlo simulation (1,000 iterations per possible subject number). The Institutional Review Board at New York University approved the experimental procedures, and all participants provided informed consent before they started the experiment.

#### Stimuli and apparatus

The target Gabor (diameter = 4°, 5 cpd, 1.25° full-width at half maximum) was presented on the left, right, upper and lower cardinal meridian locations (8° from the center to center). There were four placeholders (length = 0.16°, width = 0.06°) 0.5° away from the Gabor’s edge. The fixation cross consisted of a plus sign (length = 0.25°; width = 0.06°) at the center of the screen. The endogenous attentional cue (length = 0.75°; width = 0.2°) was presented at the center.

Participants were in a dimly lit, sound-attenuated room, with their head placed on a chinrest 57 cm away from the monitor. All stimuli were generated using MATLAB (MathWorks, MA, USA) and the Psychophysics Toolbox (Brainard, 1997; Pelli, 1997) on a gamma-corrected 20-inch ViewSonic G220fb CRT monitor with a spatial resolution of 1,280 x 960 pixels and a refresh rate of 100 Hz. To ensure fixation, participants’ eye movements were recorded using EYELINK 1000 (SR Research, Osgoode, Ontario, Canada) with a sample rate of 1,000 Hz.

#### Experimental design and procedures

Figure 2 shows the procedure of titration and the endogenous attention task. In the adapted condition, at the beginning of each block, participants adapted to a vertical 5-cpd Gabor patch flickering at 7.5 Hz in a counterphase manner, presented at the target location for 60 seconds. Each trial started with a 2s top-up phase to ensure a continuous adaptation effect throughout the block. In the non-adaptation condition, participants maintained fixation at the center for 4s (without Gabor) at the beginning of each block and for 2s at the beginning of each trial.

**Figure 2.**
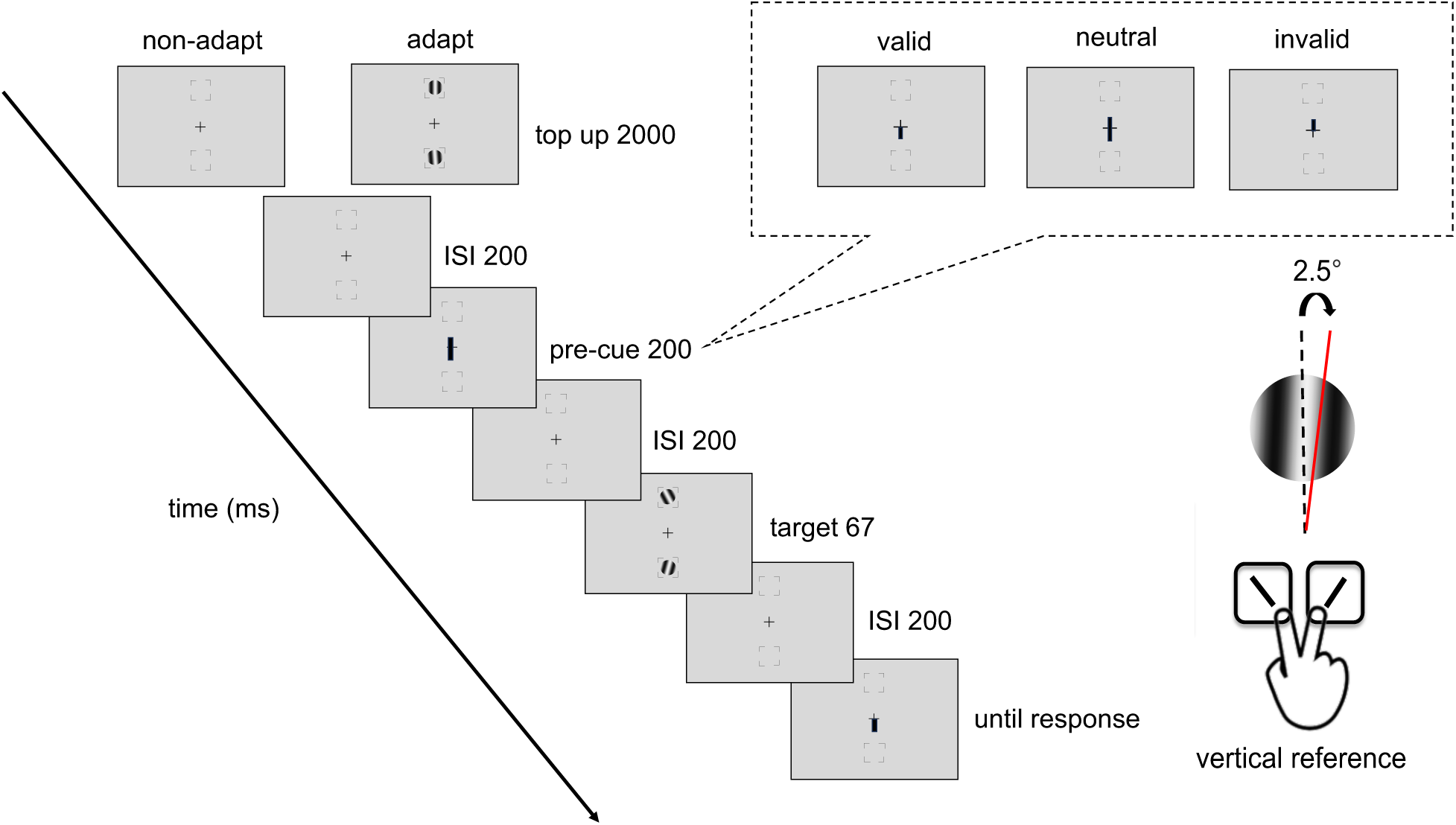
Experimental procedure: Participants performed either adaptation or nonadaptation blocks, each in separate experimental sessions. The target Gabor stimulus was always presented within the black placeholder, and target meridians were blocked. The target, two vertical Gabor stimuli were presented 8° away from the center (e.g., at the vertical meridian here; at the horizontal meridian in a different block). Participants were instructed to respond whether the Gabor was tilted clockwise or counterclockwise from vertical. The pre-cue either matches (valid condition), mismatches (invalid condition) the response cue, or does not provide location information (neutral condition). For illustration purposes, the stimulus size and spatial frequency shown here are not to scale.

After the top-up, there was a 200 ms ISI before an endogenous pre-cue was presented for 100 ms. Following a 200-ms ISI the tilted Gabor was then presented for 67 ms, followed by another 200-ms ISI and then the response cue. In a valid trial, the location indicated by the response cue matches the precue; in an invalid trial, they mismatch; in a neutral cue condition, the pre-cue points at both locations. Participants had to judge whether the target Gabor was tilted clockwise or counterclockwise off vertical. The tilt angle was 2.5°, based on pilot data and our previous study (Lee & Carrasco, 2025), to ensure an adaptation effect while avoiding floor or ceiling performance.

A feedback tone was presented when participants gave an incorrect response. The target locations were blocked in a horizontal block or a vertical block, where the target locations were presented at the horizontal or vertical meridians, respectively. Participants were asked to respond as accurately as possible while fixating at the center of the screen throughout the trial. A trial would be interrupted and repeated at the end of the block if participants’ eyes position deviated ≥1.5° from the center, from the pre-cue onset until the response cue onset.

Participants completed the adapted and non-adapted attentional task on the vertical and the horizontal meridian on different days, with a counterbalanced order. The order of horizontal and vertical meridian blocks was randomized, and the adaptation and non-adaptation titration were implemented on different days, with a counterbalanced order. There were 4 independent staircases for each adaptation condition and location, varying Gabor contrast from 2% to 85% to reach ∼75% accuracy for the orientation discrimination task. Each staircase started from 4 different points (85%, 2%, the median contrast of 43.5%, and a random point between 2% and 85%) and contained 48 trials. Four blocks (192 trials per location for each adaptation and non-adaptation conditions) were conducted consecutively for the horizontal meridian block or the vertical meridian block. The contrast threshold was derived using an adaptive staircase procedure using the Palamedes toolbox (Prins & Kingdom, 2018), as in previous studies (e.g., Fernández & Carrasco, 2020; Hanning et al., 2022; Jigo & Carrasco, 2018; Lee & Carrasco, 2025; Lee et al., 2024) and averaging the last 8 trials. The Gabors were always preceded by a neutral pre-cue, which, as in many studies (e.g., Dosher & Lu, 2000b; Fernández et al., 2022; Huang, Liao, Chen, & Chen, 2025; Jigo & Carrasco, 2020; Li, Pan, & Carrasco, 2021; Luzardo & Yeshurun, 2025; Palmieri & Carrasco, 2024; Ramamurthy, White, & Yeatman, 2024; Tünçok, Carrasco, & Winawer, 2025), provided the same temporal information as the valid and invalid cues, but no information about the spatial location.

In this endogenous attention task, for each adapted and non-adapted condition, 20% of the trials had a neutral cue, which pointed at both locations; 80% of the trials had an attentional cue pointing toward a location, 75% among them were valid cues, and the other 25% were invalid cues. All participants completed a practice session to familiarize themselves with the task procedure.

#### Psychometric function fitting

We fitted a Weibull function for the accuracy as a function of contrast threshold. For each location and adaptation condition, a logistic function was fit to the data using maximum likelihood estimation using the fmincon function in MATLAB. The results derived from the psychometric function estimation positively correlated (*ps<*.01) with the staircase results in all experiments, verifying our procedure in all conditions.

#### Behavioral data analyses

Behavioral data analyses were performed using R (Team, 2000). A three-way repeated-measures analysis of variance (ANOVA) on d’ was conducted on the factors of location (horizontal meridian, upper, lower), adaptation (adapted, non-adapted), and attention (valid, neutral, invalid) conditions to assess statistical significance. Repeated-measures ANOVA along with effect size (*η^2^*) were computed in R and used to assess statistical significance.

## Results

### Adaptation effect varied around polar angle

After deriving the c50 contrast for the horizontal meridian (HM), upper, and lower vertical meridians for both the adapted and non-adapted conditions, we conducted a two-way ANOVA on contrast thresholds (Figure 3). This analysis showed a main effect of location [*F*(2,22)=7.89, *p*=.003, *ηp^2^*=0.42] and a higher threshold in the adapted than non-adapted conditions [*F*(1,11)=18.44, *p*=.001, *ηp^2^*=0.63], and an interaction [*F*(2,22)=3.58, *p*=.045, *ηp^2^*=0.25], indicating that the adaptation effect varied across locations.

**Figure 3.**
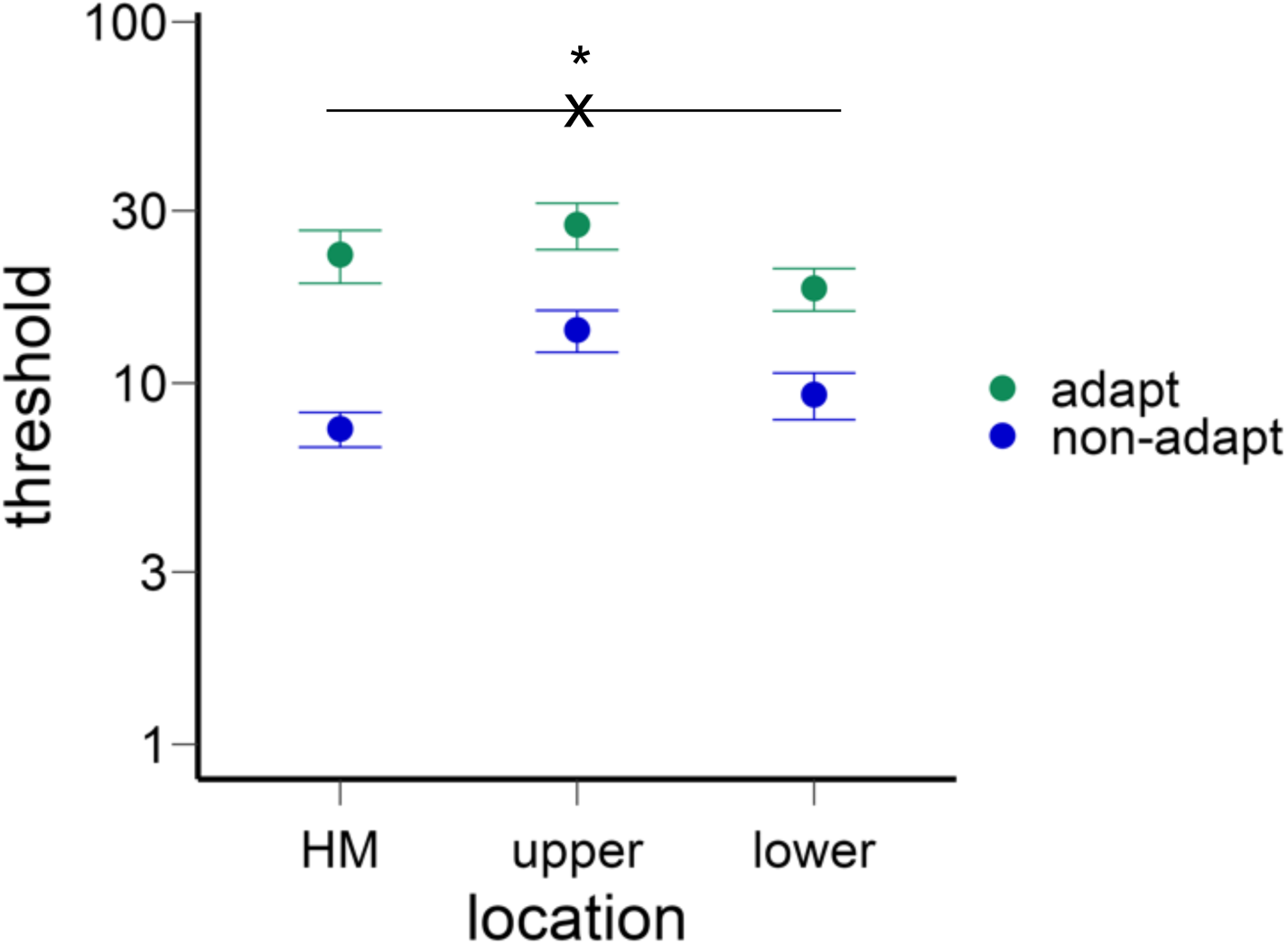
The contrast thresholds for different locations and adaptation conditions. The thresholds were higher in the vertical than horizontal meridian (HM), and higher in the upper than lower vertical meridian. The thresholds were also higher in the adapted than non-adapted conditions. Critically, the adaptation effect was stronger in the horizontal than vertical meridian. The error bars indicate ±1 SEM.

We confirmed that the HVA and VMA emerged in the non-adaptation condition (Figure 3): Contrast thresholds were lower along the horizontal than the vertical meridian [*t*(11)=5.87, *p*<.001, *d*=1.69) and lower at the lower than upper vertical meridian [*t*(11)=2.37, *p*=.037, *d*=0.68].

Next, we assessed the adaptation effect at the horizontal and vertical meridians. The normalized adaptation effect (calculated as the difference between adapted and non-adapted thresholds divided by the sum of the thresholds, as in Lee and Carrasco (2025) was stronger at the horizontal than the vertical meridian [*t*(11)=3.39, *p*=.006, *d*=0.98] (Figure 3, see gaps between adapt and non-adapt conditions for different locations), but no significant difference between the upper and lower vertical meridian [*t*(11)<1].

### Endogenous attentional effect

Figure 4 shows the results. We compared the endogenous attentional effect on d′ by conducting a three-way ANOVA on the factors of location (HM, upper, lower), attentional validity (valid, neutral, invalid), and adaptation (adaptation, non-adaptation). Given that we titrated the contrast thresholds across locations and adaptation conditions, we expected no main effects of either adaptation or location. Indeed, there was a main effect of attention [*F*(2,22)=53.18, *p*<.001, *ηp^2^*=0.83], but neither of location [*F*(2,22)<1], nor of adaptation [*F*(1,11)<1]. There was neither a 3-way interaction nor 2-way interactions [all *ps*>.1].

**Figure 4.**
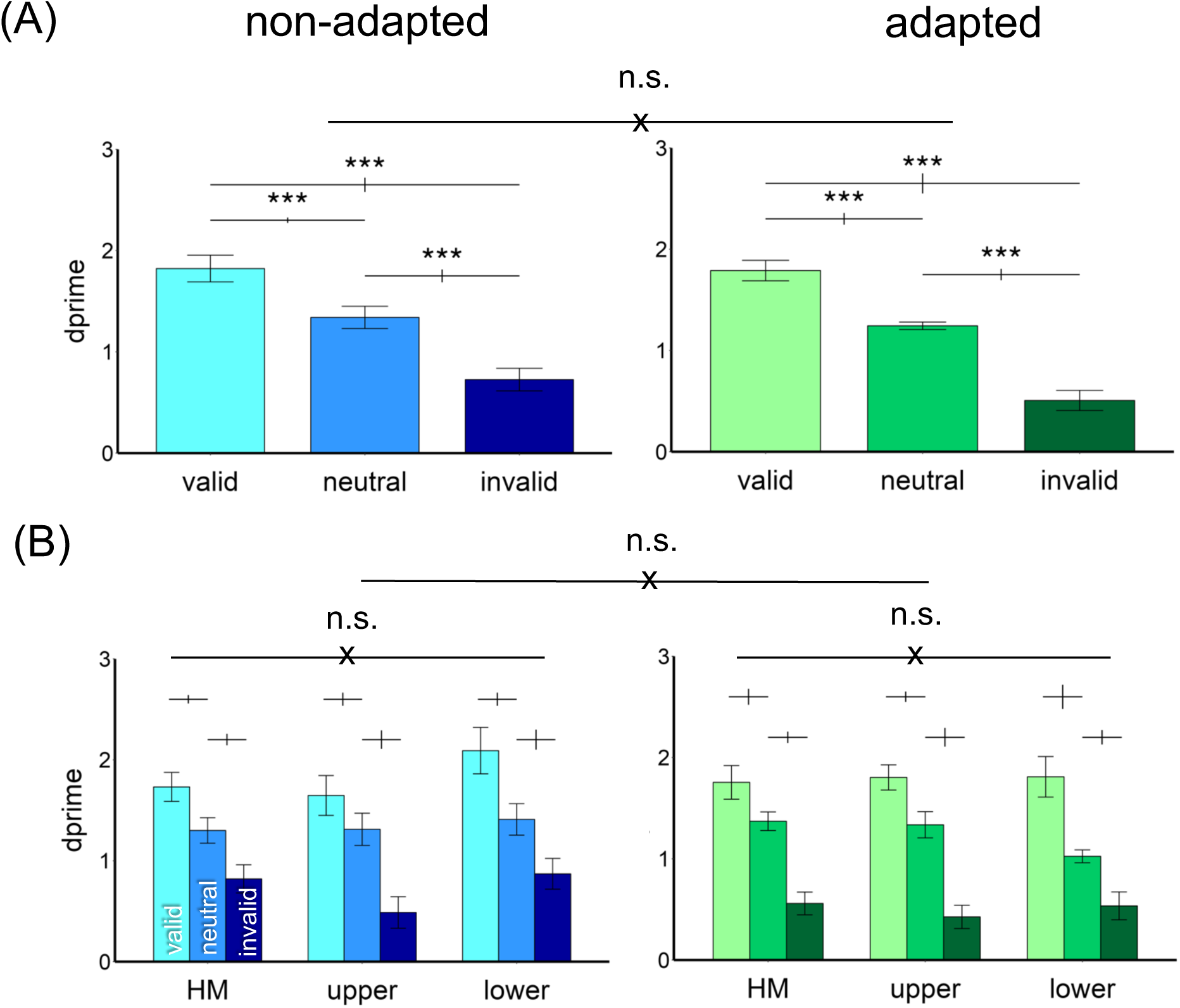
Performance in Experiment 1. (A) d′ was higher in the valid followed by neutral and invalid conditions in both non-adapted and adapted conditions. There was no difference between the adapted and non-adapted conditions. (B) The attentional effects were similar around polar angle –horizontal meridian (HM), and upper and lower vertical meridian– and were comparable in the adapted and non-adapted conditions. The error bars above the bar plots indicate ±1 SEM of the difference between conditions. *** p<.001, n.s. p>.05.

The results were further confirmed by separating the adapted and non-adapted conditions into two 2-way ANOVAs on attention and location. For the non-adapted condition, we observed a main effect of attention [*F*(2,22)=46.74, *p*<.001, *ηp^2^*=0.81] but not of location [*F*(2,22)<1] or an interaction [*F*(4,44)=1.68, *p*>.1]. The same pattern emerged for the adapted condition: a main effect of attention [*F*(2,22)=38.59, *p*<.001, *ηp^2^*=0.78] but not of location [*F*(2,22)<1] or an interaction [*F*(4,44)=1.48, *p*>.1]. Thus, neither adaptation state nor location modulated the pronounced overall effect of attention.

We plot the individual data for the endogenous attentional effect (valid *d′* − invalid *d′*) in the adapted and non-adapted conditions (Figure 5). There was no difference between the two conditions [*t*(11)=1.27, *p*>.1].

**Figure 5.**
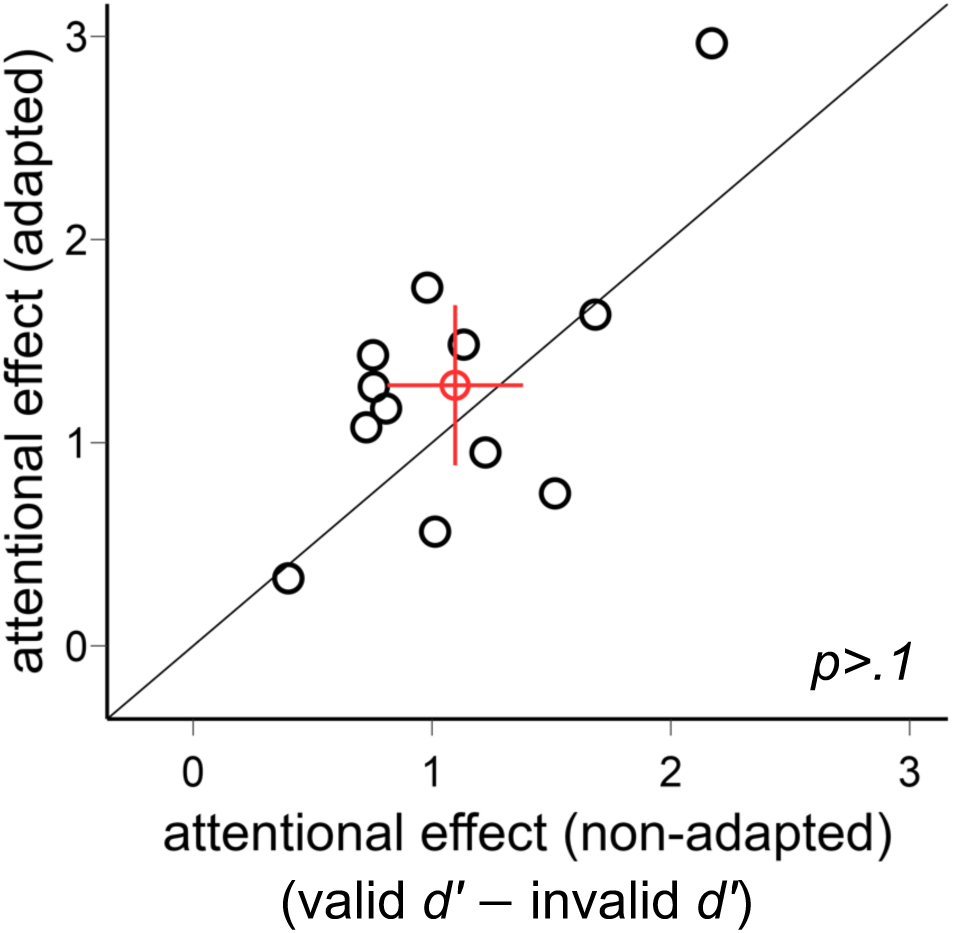
Comparison of the endogenous attentional effects (valid d′ − invalid d′) in Experiment 1. The attentional effects were comparable in the adapted and non-adapted conditions. The red circle indicates the mean of all participants, and the error bars indicate ±1 SEM.

In sum, the endogenous attentional effect was comparable across locations and adaptation conditions.

### Experiment 2 — Exogenous attention

Experiment 1 shows that endogenous attention does not reshape the performance fields, even after differential adaptation effects across meridians. In Experiment 2, we examined whether exogenous attention exhibits a similar or distinct pattern as endogenous attention, given their well-established differences in temporal dynamics: Whereas endogenous attention takes about 300 ms to deploy and its effects can be sustained for many seconds, exogenous attention effects peak at about 120 ms and its effects are transient (reviews: Carrasco, 2011, 2014; Carrasco & Barbot, 2014). Moreover, endogenous attention is flexible whereas exogenous attention is not (e.g., Barbot & Carrasco, 2017; Barbot et al., 2012; Giordano et al., 2009; Hein et al., 2006; Nakayama & Mackeben, 1989; Yantis & Jonides, 1996; Yeshurun & Carrasco, 1998, 2008), and the effects of endogenous attention scale with cue validity, whereas those of exogenous attention do not (e.g., Giordano et al., 2009; Kinchla, 1980; Mangun & Hillyard, 1990; Sperling & Melchner, 1978).

To manipulate exogenous attention, we used a peripheral cue (a bolded placeholder) presented before the target onset. According to a normalization model of attention, exogenous attention can also affect contrast gain when the attentional window is large enough (Reynolds & Heeger, 2009; Herrmann et al., 2010). To induce a large attentional window while maintaining overlap between the target and adaptors and ensure the adaptation effect, we randomly presented the target in one of the five locations within the placeholders (Figure 6), and participants were explicitly instructed to attend to the whole space encompassed by the placeholder, as the target could appear anywhere within the placeholder. This procedure has been successfully used to manipulate the size of the attentional window in both exogenous and endogenous covert spatial attention, as well as in presaccadic attention (e.g., Binda & Murray, 2015; Cutrone, Heeger, & Carrasco, 2018; Feng & Spence, 2017; Grubb et al., 2013; Herrmann et al., 2010; Li et al., 2021).

**Figure 6.**
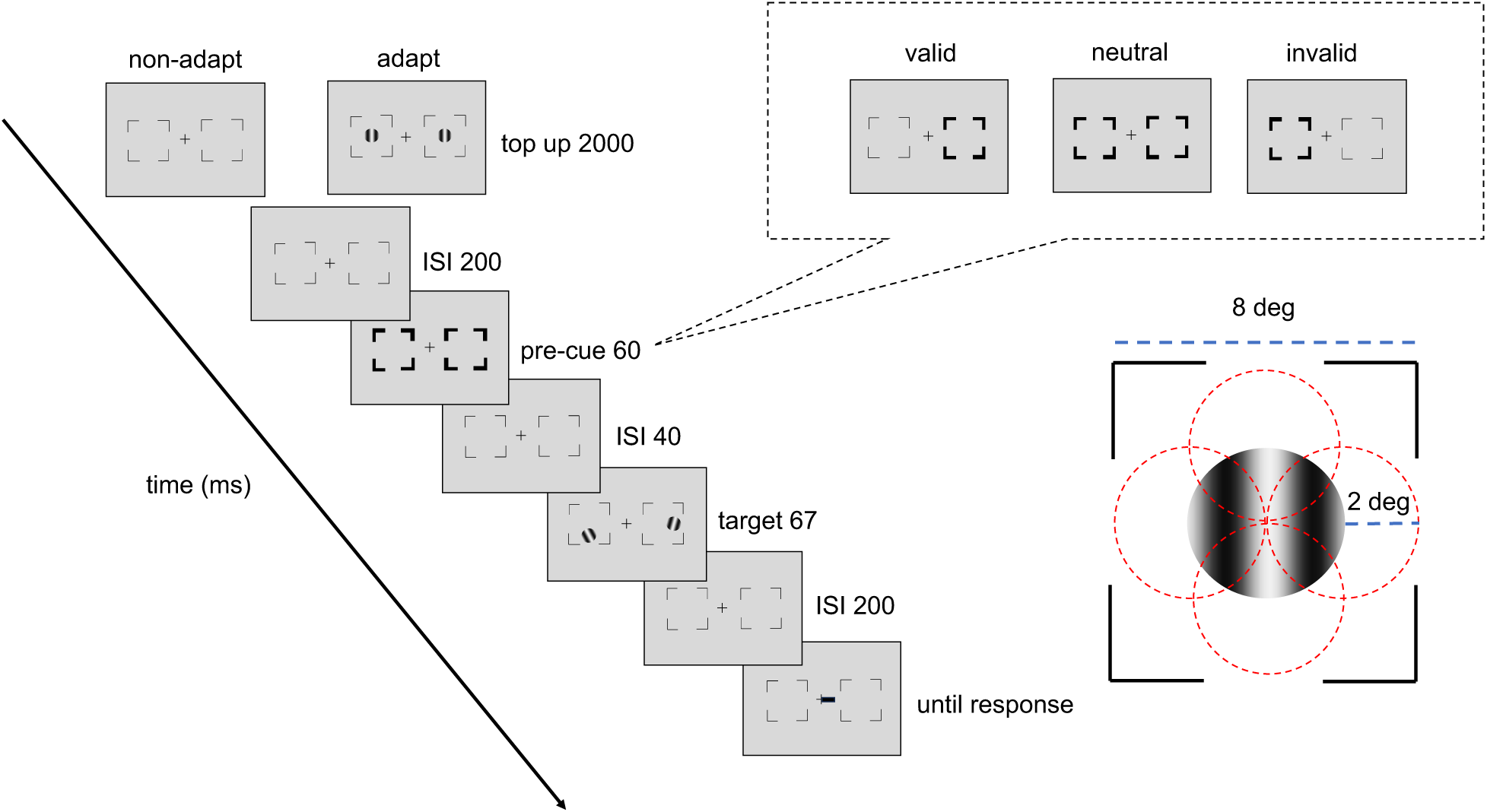
Experimental procedure: The procedure was the same as Experiment 1 except for the pre-cue and ISI timings. The pre-cue (bolded placeholders) either matches (valid condition), mismatches (invalid condition) the response cue, or does not provide location information (neutral condition). The placeholders were wider (8°) than Experiment 1. The target, two vertical Gabor stimuli were presented on average 8° away from the center (e.g., at the horizontal meridian for example here; at the vertical meridian in a different block). There were 5 possible target locations, which were 2° away from the central Gabor. For illustration purposes, the stimulus size and spatial frequency shown here are not to scale.

## Methods

### Participants

Eleven out of 12 participants^1^ who participated in Experiment 1, including author HHL, also participated in Experiment 2. We tested the same group of participants to compare the results from endogenous and exogenous attentional effects after adaptation.

### Stimuli and apparatus

Figure 6 shows an experimental trial. The target stimuli and the apparatus were the same as Experiment 1. The placeholders in Experiment 2 (length = 0.256° for placeholders that were further away from the center, length = 0.192° for placeholders that were closer to the center, all width = 0.06°) were larger, given that there were 5 possible target locations: center and 2° on the upper, lower, left, or right of the central Gabor. During the cue presentation, the placeholders became thicker (6 pixels bigger for the frame elements closer to the center and 8 pixels bigger for the frame elements further away from the center) to capture participants’ exogenous attention.

### Experimental design and procedures

The same c50 contrast derived from Experiment 1 was used in Experiment 2 for the adapted and non-adapted conditions across locations. The experimental design and procedures were the same as in Experiment 1, except for the following: After the top-up, there was a 200-ms ISI before the exogenous pre-cue appeared for 60 ms, followed by 40-ms ISI. The tilted Gabor was then presented for 67 ms followed by another 200-ms ISI and the response cue. Participants were explicitly told that the exogenous cues were not informative; i.e., they were equally likely to be valid, neutral and invalid (33% each). Participants were instructed to enlarge their attentional window during the task, as they were explicitly told that the target could appear anywhere within the placeholders.

## Results

Figure 7 shows our results. As in Experiment 1, we compared the exogenous attentional effect on d′ by conducting a three-way ANOVA on the factors of location (HM, upper, lower), attentional validity (valid, neutral, invalid), and adaptation (adaptation, non-adaptation). There was a main effect of attention [*F*(2,20)=20.7, *p*<.001, *ηp^2^*=0.67], but neither of location [*F*(2,20)=2.61, *p*=.099], nor of adaptation [*F*(1,10)=1.08, *p*>.1]. There was neither a 3-way interaction [*F*(4,40)=2.36, *p*=.069] nor 2-way interactions [all *ps*>.1].

**Figure 7.**
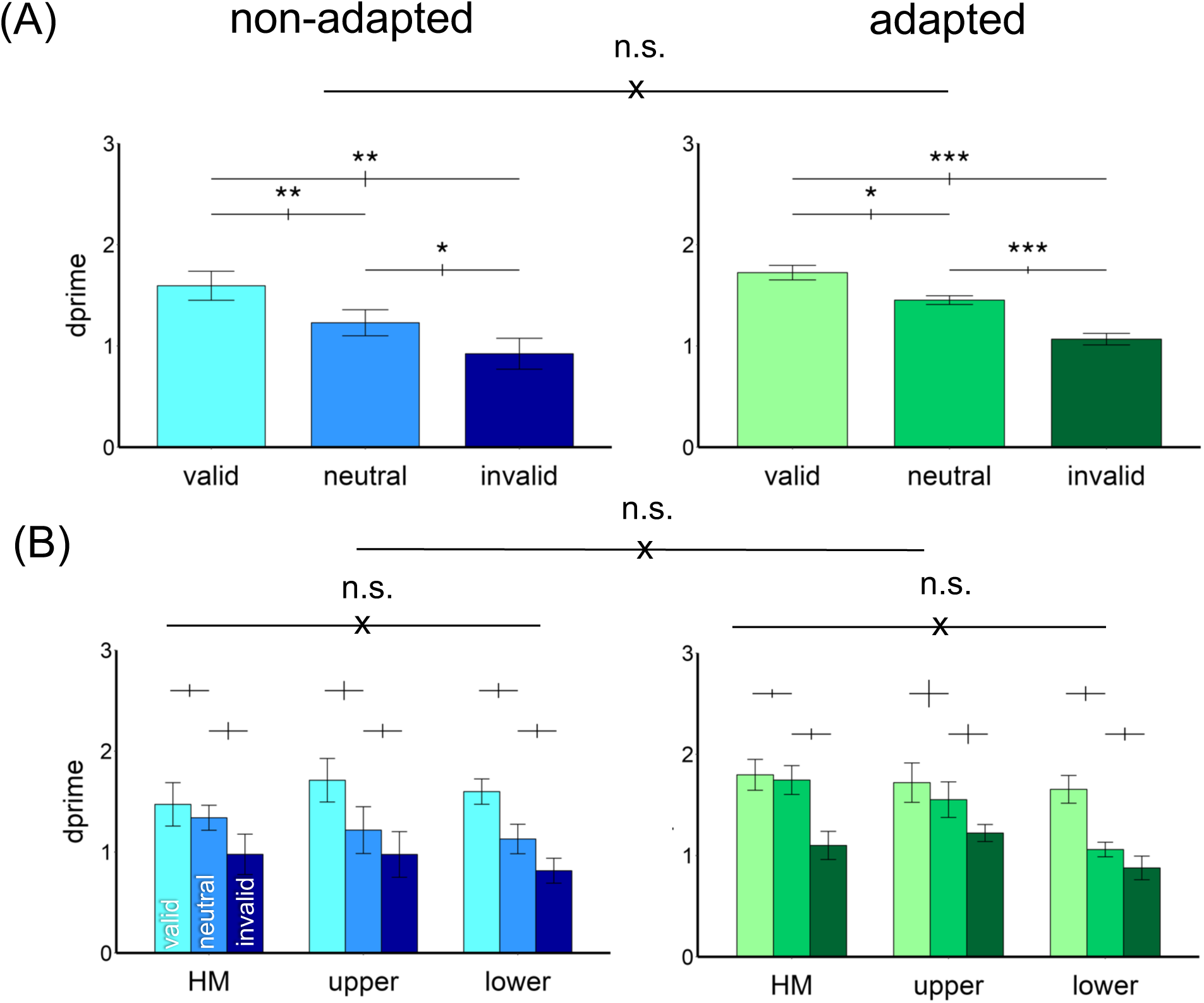
Performance in Experiment 2. (A) d′ was higher in the valid followed by neutral and invalid conditions in both non-adapted and adapted conditions. There was no difference between the adapted and non-adapted conditions. (B) The attentional effects were similar around polar angle –horizontal meridian (HM), and upper and lower vertical meridian– and were comparable in the adapted and non-adapted conditions. The error bars above the bar plots indicate ±1 SEM of the difference between conditions. *** p<.001, ** p<.01, * p<.05, n.s. p>.05.

The results here were further confirmed by separating the adapted and non-adapted conditions into two 2-way ANOVAs on attention and location. For the non-adapted condition, we observed a main effect of attention [*F*(2,20)=13.33, *p*<.001, *ηp^2^*=0.57] but neither of location [*F*(2,20)<1], nor an interaction [*F*(4,40)=1.33, *p*>.1]. The same pattern emerged for the adapted condition: a main effect of attention [*F*(2,20)=21.19, *p*<.001, *ηp^2^*=0.68] but neither of location [*F*(2,20)=2.33, *p*>.1] nor an interaction [*F*(4,40)=2.01, *p*>.1].

We plot the individual data for the exogenous attentional effect (valid *d′* − invalid *d′*) in the adapted and non-adapted conditions (Figure 8). There was no difference between the two conditions [*t*(10)<1].

**Figure 8.**
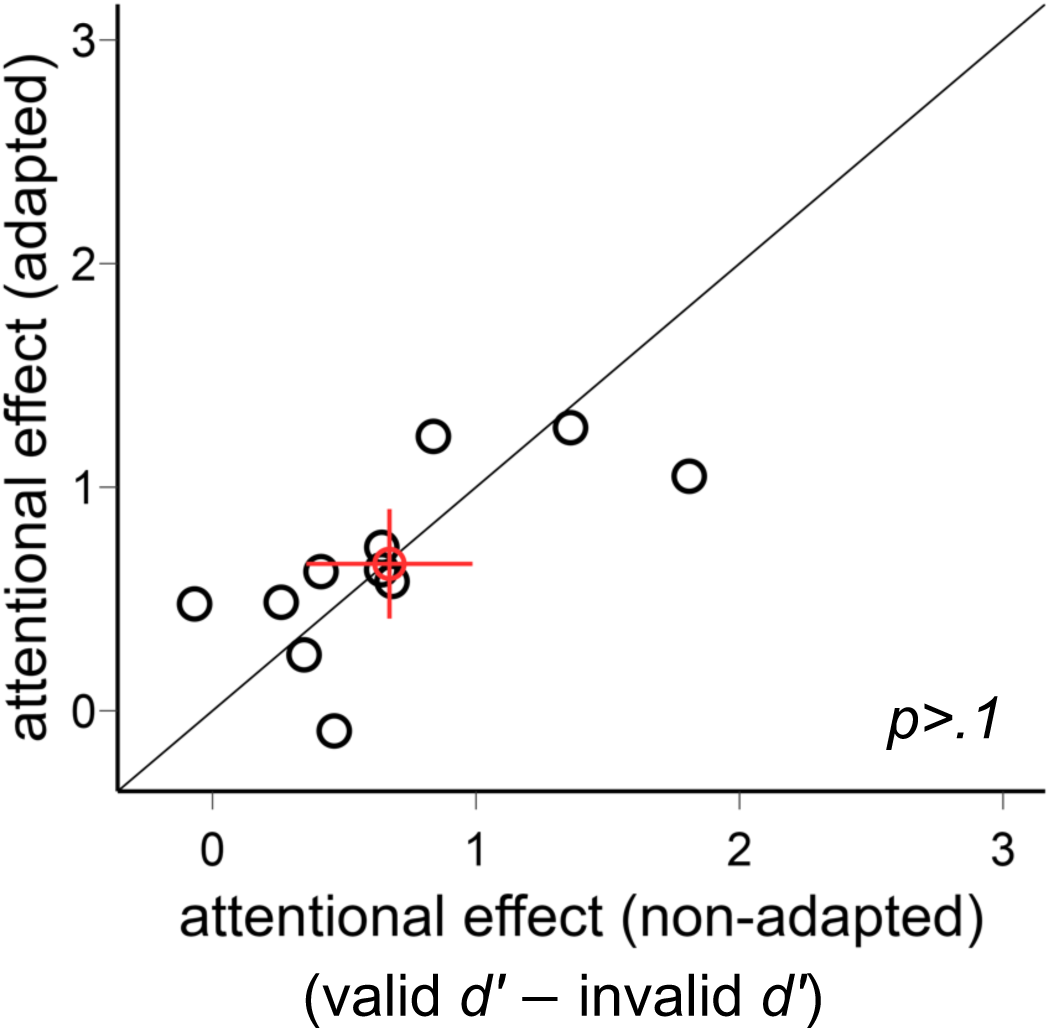
Comparison of the exogenous attentional effects (valid d′ − invalid d′) in Experiment 2. The attentional effects were comparable in the adapted and non-adapted conditions. The red circle indicates the mean of all participants, and the error bars indicate ±1 SEM.

In sum, the exogenous attentional effect was comparable across locations and adaptation conditions.

### Comparing Experiments 1 and 2

Given that we had 11 common participants in Experiments 1 and 2, we conducted a 4-way repeated-measures within-subject ANOVA on the factor of type of attention (endogenous, exogenous), attentional validity (valid, neutral, invalid), adaptation (adapted, non-adapted), and location (HM, upper, lower). There was a main effect of attentional validity [*F*(2,20)=51.72, *p*<.001, *ηp^2^*=0.84], and an interaction between attentional validity and type of attention [*F*(2,20)=7.38, *p*=.004, *ηp^2^*=0.42]. Post-hoc analyses indicated that valid condition had the highest *d′* followed by neutral [valid – neutral: *t*(10)=6.78, *p*<.001, *d*=2.04] and invalid conditions [neutral – invalid: *t*(10)=6.17, *p*<.001, *d*=1.86]. The attentional effect (valid *d′* – invalid *d′*) was stronger for endogenous than exogenous attention [*t*(10)=2.95, *p*=.015, *d*=0.89]. Importantly, there were no 4-way interaction [*F*(4,40)<1] nor any other significant effects [all *ps*>.05], indicating that the effect for both types of attention did not vary across locations nor across adaptation conditions.

Furthermore, we found a positive Pearson correlation [*r*=.39, *p*=.025] between the exogenous and endogenous overall attentional effect (collapsing across adaptation conditions and locations), which indicates that those observers who had a stronger effect of one type of attention also had a stronger effect for the other type.

## Discussion

In this study, we investigated whether attention interacts with adaptation around polar angle. Our results are consistent with separate studies showing: (1) without adaptation, the typical performance fields emerged, with lower contrast thresholds at the horizontal than the vertical meridian (HVA) and at the lower than the upper vertical meridian (VMA) (e.g., Abrams et al., 2012; Baldwin et al., 2012; Cameron et al., 2002; Carrasco et al., 2001; Corbett & Carrasco, 2011; Fuller et al., 2008; Himmelberg et al., 2020; Lee & Carrasco, 2025); (2) adaptation effects were stronger at the horizontal than the vertical meridian (Lee & Carrasco, 2025); and (3) both endogenous attention (Purokayastha et al., 2021; Tünçok, Carrasco, & Winawer, 2025) and exogenous attention (Cameron et al., 2002; Carrasco et al., 2001; Roberts et al., 2018; Roberts et al., 2016) enhanced contrast sensitivity similarly across all tested locations. Furthermore, our results revealed that: (1) endogenous attention restored contrast sensitivity following adaptation, (2) endogenous and exogenous attention had similar effects on contrast sensitivity before and after adaptation–both enhanced contrast sensitivity at the attended location, with concomitant costs at unattended locations, and (3) they did so uniformly at the cardinal meridians around the visual field–despite differential adaptation effects.

The finding that endogenous attention enhances contrast sensitivity to a similar extent in adapted and non-adapted conditions indicates that visual adaptation does not modulate the attentional effect. This novel finding is consistent with corresponding findings for exogenous attention on contrast sensitivity after adaptation (Lee et al., 2024; Pestilli et al., 2007). Despite its flexible nature (reviews: Carrasco, 2011, 2014; Carrasco & Barbot, 2014; Olivers, 2025), endogenous attention neither increased nor decreased contrast sensitivity differentially before and after adaptation, indicating that these two processes, which help manage limited bioenergetic resources, play independent roles in shaping performance.

Typically, the effect of exogenous attention manifests as response gain and the effect of endogenous attention as contrast gain (Ling & Carrasco, 2006a; Pestilli et al., 2009). In the exogenous attention experiment, we induced contrast gain by manipulating the size of the attentional window, presenting the target Gabor at one of 5 different locations within a larger stimulus placeholder. According to Reynolds and Heeger’s normalization model of attention, attention produces contrast gain rather than response gain when the attentional window is large relative to stimulus size (Reynolds & Heeger, 2009), a prediction confirmed psychophysically and with functional magnetic resonance imaging (fMRI) (Herrmann et al., 2010). By contrast, endogenous attention can induce response gain when deployed over a relatively smaller attentional window than the stimulus size (Fernández et al., 2023; Herrmann et al., 2010; Morrone, Denti, & Spinelli, 2004). Consistent with previous findings (Lee et al., 2024; Pestilli et al., 2007), exogenous attention modulated contrast sensitivity to a similar extent in adapted and non-adapted conditions, indicating that adaptation did not modulate its effect. These results support Hypothesis 1: after adaptation, covert spatial attention modulates contrast sensitivity to the same extent as without adaptation (Figure 1A).

In this exogenous attention experiment, to induce a larger attentional window, participants were explicitly told that the target could appear anywhere within the placeholders. This manipulation should not affect the effects of exogenous attention as it cannot induce endogenous attention. The effects of endogenous attention scale with cue validity (e.g., Giordano et al., 2009; Kinchla, 1980; Mangun & Hillyard, 1990; Sperling & Melchner, 1978), and in the exogenous attention experiment, the cue was uninformative: each of the valid, invalid, and neutral cues was presented on 33% of the trials, so when a cue indicated one location out of two, its validity was 50%. Thus, had observers deployed endogenous attention in Experiment 2, performance would have been similar for valid and invalid conditions. Instead, we found significant benefits at the attended location and significant costs at unattended locations, consistent with an exogenous attention effect. Moreover, given the timing of the exogenous cue (∼120 ms)–and that endogenous attention takes ∼300 ms to be deployed (e.g., Cheal, Lyon, & Hubbard, 1991; Geweke, Pokta, & Störmer, 2021; Liu, Stevens, & Carrasco, 2007; Nakayama & Mackeben, 1989; Remington, Johnston, & Yantis, 1992, for reviews, see Carrasco 2011, 2014, Carrasco & Barbot, 2015)–endogenous attention could not contribute.

Adaptation was more pronounced at the horizontal than the vertical meridian. Unlike our previous study (Lee & Carrasco, 2025), which blocked each target location, here we introduced greater target uncertainty by using two possible target locations per trial (Figure 2). The replication of the adaptation pattern across studies shows that that the previous findings are robust to target uncertainty and generalize across participants. Most adaptation studies have examined only the horizontal meridian (e.g., Beaton & Blakemore, 1981; Carrasco et al., 2006; Gao, Webster, & Jiang, 2019; Greenlee, Georgeson, Magnussen, & Harris, 1991; Pestilli et al., 2007; Schieting & Spillmann, 1987), only the vertical meridian (e.g., Bell, Gheorghiu, Hess, & Kingdom, 2011; Bell, Gheorghiu, & Kingdom, 2009; Montaser-Kouhsari & Rajimehr, 2004), or did not analyze target locations separately (e.g., Bao, Fast, Mesik, & Engel, 2013; Lin, Zhou, Naya, Gardner, & Sun, 2021; Ling & Carrasco, 2006b; Zimmermann, Weidner, Abdollahi, & Fink, 2016). Our results add further evidence that adaptation differs across meridians, an important finding to consider in future studies and models of vision.

Endogenous and exogenous attention enhanced contrast sensitivity similarly around polar angle, despite the differential effects of adaptation. Consistent with previous studies (Cameron et al., 2002; Carrasco et al., 2001; Roberts et al., 2018; Roberts et al., 2016), asymmetries at the cardinal locations were resistant to both endogenous and exogenous attention, indicating their resilient nature and that they cannot be easily reshaped. In contrast, consistent with a recent finding (Lee & Carrasco, 2025), visual adaptation reduced contrast sensitivity more at the horizontal than the vertical meridian, yet neither type of covert spatial attention modulated the extent of the asymmetries altering the shape of the performance fields, notwithstanding the differential adaptation effect. This similar effect is notable given that endogenous attention is flexible and exogenous attention automatic (e.g., Carrasco, 2011, 2014; Carrasco & Barbot, 2014; Olivers, 2025), yet neither compensated for poor performance. These findings provide further evidence regarding the resilience of polar angle asymmetries and support Hypothesis 4 (Figure 1D): visual adaptation does not modulate the effects of covert spatial attention, even at the location of poorest performance.

What contributes to performance asymmetries in the HVA and VMA? These asymmetries arise from both retinal and cortical factors. Retinally, cone density is higher at the horizontal than the vertical meridian (Curcio, Sloan Jr, Packer, Hendrickson, & Kalina, 1987; Curcio, Sloan, Kalina, & Hendrickson, 1990), and midget-RGC density is higher at the lower than the upper vertical meridian (Curcio et al., 1990; Song, Chui, Zhong, Elsner, & Burns, 2011). Cortically, V1 surface area is larger for the horizontal than the vertical meridian, and for the lower than the upper vertical meridian (Benson, Kupers, Barbot, Carrasco, & Winawer, 2021; Himmelberg et al., 2021; Himmelberg, Kwak, Carrasco, & Winawer, 2025; Himmelberg, Tünçok, et al., 2023; Himmelberg, Winawer, & Carrasco, 2022, 2023; Lee & Carrasco, 2025; Silva et al., 2018). Moreover, cortical factors account for more variance in these asymmetries than retinal factors (Kupers, Benson, Carrasco, & Winawer, 2022). Still, these factors cannot fully explain behavioral differences observed in psychophysical tasks, which are diminished but still present once stimulus size is cortically magnified (Jigo, Tavdy, Himmelberg, & Carrasco, 2023), suggesting that additional factors –such as sensory tuning and neuronal computations– also contribute to the HVA and VMA (Himmelberg, Winawer, & Carrasco, 2023; Jigo et al., 2023; Xue, Barbot, Abrams, Chen, & Carrasco, 2025).

Endogenous and exogenous attention rely on different neural substrates. fMRI studies show differential activity modulation across the frontoparietal network (Beck & Kastner, 2014; Buschman & Miller, 2009; Chica, Bartolomeo, & Lupiáñez, 2013; Fiebelkorn & Kastner, 2020; Kastner & Buschman, 2017; Meyer, Du, Parks, & Hopfinger, 2018), temporoparietal junction (Dugué, Merriam, Heeger, & Carrasco, 2018), and visual cortex (Dugué, Merriam, Heeger, & Carrasco, 2020; Hopfinger & West, 2006; Ling, Jehee, & Pestilli, 2015). Transcranial magnetic stimulation (TMS) studies, which disrupt the neuronal balance between excitation and inhibition (Bradley, Nydam, Dux, & Mattingley, 2022; Kobayashi & Pascual-Leone, 2003; Valero-Cabré, Pascual-Leone, & Coubard, 2011), revealed that early visual cortex plays a critical role for adaptation (Lee et al., 2024; Perini et al., 2012) and exogenous attention (Fernández & Carrasco, 2020; Lee et al., 2024), whereas the human homologue of the right frontal eye fields (rFEF+) plays a critical role for endogenous attention (Fernández et al., 2023). Critically, disrupting rFEF+ does not affect exogenous attention (Chen et al., 2025), and disrupting early visual cortex does not affect endogenous attention (Fernández et al., 2023), indicating a double dissociation. Despite these distinct neural underpinnings, both types of covert spatial attention affected contrast sensitivity uniformly at the cardinal meridians around polar angle and did not interact with location or adaptation. These findings suggest that distinct neuronal populations underlie polar angle asymmetries, adaptation, and attentional modulation.

We found stronger attentional effects for endogenous than exogenous attention. Given that adaptation is more effective when the adaptor and the target spatially overlap (Kovács, Zimmer, Harza, & Vidnyánszky, 2007; Larsson & Harrison, 2015; Webster, 2011, 2015), we introduced target uncertainty with 5 possible target locations and allowed 2° overlap between adaptor and target to elicit adaptation while allowing exogenous attention to operate via contrast gain. This manipulation may have yielded a slightly narrower exogenous attentional window than for endogenous attention in our design, as well as compared with previous studies. For example, Herrmann et al. (2010) used five possible target locations with no overlap, whereas in our current study the target Gabors could overlap by 2° within placeholders. According to Reynolds and Heeger’s normalization model of attention (Reynolds & Heeger, 2009), attention multiplies stimulus-evoked activity before divisive normalization. In our task, normalization may have pooled a broader suppressive drive than in typical exogenous attention tasks, but not as broad as in typical endogenous attention tasks —leading to less pronounced contrast gain and thus weaker exogenous than endogenous attention effects.

Why do type of spatial covert attention, adaptation, and polar angle asymmetries not interact? The visual cortex plays a crucial role in all three processes. fMRI studies have shown that covert endogenous spatial attention modulates activity in visual cortex via feedback from frontoparietal cortex (Buschman & Miller, 2009; Chica et al., 2013; Corbetta, Patel, & Shulman, 2008; Corbetta & Shulman, 2002; Dugué et al., 2020; Lauritzen, D’Esposito, Heeger, & Silver, 2009; Pestilli, Carrasco, Heeger, & Gardner, 2011) and increasingly modulates activity in the occipital visual areas (Dugué et al., 2020), with V1/V2, its early visual areas, being not critical for endogenous attention, as TMS on these areas does not alter its effect on visual perception (Fernández et al., 2023). In contrast, exogenous attention modulates visual cortex via feedforward activation (Dassanayake, Michie, & Fulham, 2016; Dugué et al., 2020; Hopfinger, Luck, & Hillyard, 2004; Liu, Pestilli, & Carrasco, 2005; F. Wang, Chen, Yan, Zhaoping, & Li, 2015; Westerberg, Schall, Woodman, & Maier, 2023), and V1/V2 are critical for its effect, as TMS on these areas eliminates the effect of exogenous attention on visual perception (Fernández & Carrasco, 2020; Lee et al., 2024). Moreover, these two attention types also differentially modulate visual subregions of the temporoparietal-junction (Dugué et al., 2018). All these differences underscore the distinct contributions of endogenous and exogenous in modulating visual perception.

Early visual cortex also plays a critical role in visual adaptation. TMS over V1/V2 decreases contrast adaptation (Perini et al., 2012), and adaptation modulates contrast response functions in V1/V2 (Altan et al., 2025; Gardner et al., 2005; Vinke, Bloem, & Ling, 2022). A TMS study revealed that adaptation and exogenous attention interact in early visual cortex (Lee et al., 2024), but it is unknown whether they do so systematically around polar angle, as several factors shape asymmetries. Moreover, it is presently unknown whether endogenous attention and adaptation interact either in early occipital or in frontal areas.

Both adaptation (Lee & Carrasco, 2025) and polar angle asymmetries (Benson et al., 2021; Himmelberg et al., 2022; Himmelberg, Winawer, & Carrasco, 2023; Lee & Carrasco, 2025) correlate with V1 surface area, but surface area alone cannot fully account for these asymmetries (Jigo et al., 2023). Additional factors such as neural gain also contribute to these asymmetries (Xue et al., 2025). Future research integrating computational modeling, neuroimaging, neurostimulation, and psychophysics will be essential to assess the relative contributions of cortical and computational factors to attention, adaptation and polar angle asymmetries.

In conclusion, this study reveals that performance asymmetries are resistant to the effects of both endogenous and exogenous covert spatial attention, despite their distinct temporal dynamics and differences in flexibility–even after adaptation induces differential effects across meridians. Although both adaptation and attention help allocate limited resources according to task demands, neither type of covert spatial attention differentially enhance target processing at locations that differ in intrinsic discriminability and their corresponding representation in cortical surface area.

## Acknowledgments

This study was supported by NIH NEI R01-EY027401 to M.C. and the Ministry of Education in Taiwan and 2025 National Science and Technology Council (NSTC) Taiwanese Overseas Pioneers Grants (TOP Grants) for PhD Candidates to H.-H.L. We thank Nick Crotty, David Tu and Shutian Xue for their feedback on a previous version of the manuscript.

## Footnotes

1 One participant was no longer around NYC to participate

## References

1. Abrams, J., Nizam, A., & Carrasco, M. (2012). Isoeccentric locations are not equivalent: The extent of the vertical meridian asymmetry. Vision Research, 52(1), 70–78.

2. Afraz, A., Pashkam, M. V., & Cavanagh, P. (2010). Spatial heterogeneity in the perception of face and form attributes. Current Biology, 20(23), 2112–2116.

3. Altan, E., Morgan, C. A., Dakin, S. C., & Schwarzkopf, D. S. (2025). Spatial frequency adaptation modulates population receptive field sizes. elife, 13, RP100734.

4. Altpeter, E., Mackeben, M., & Trauzettel-Klosinski, S. (2000). The importance of sustained attention for patients with maculopathies. Vision Research, 40(10-12), 1539–1547.

5. Baldwin, A. S., Meese, T. S., & Baker, D. H. (2012). The attenuation surface for contrast sensitivity has the form of a witch’s hat within the central visual field. Journal of Vision, 12(11), 23.

6. Bao, M., Fast, E., Mesik, J., & Engel, S. (2013). Distinct mechanisms control contrast adaptation over different timescales. Journal of Vision, 13(10), 14.

7. Barbot, A., & Carrasco, M. (2017). Attention modifies spatial resolution according to task demands. Psychological Science, 28(3), 285–296.

8. Barbot, A., Landy, M. S., & Carrasco, M. (2012). Differential effects of exogenous and endogenous attention on second-order texture contrast sensitivity. Journal of Vision, 12(8), 6.

9. Barbot, A., Xue, S., & Carrasco, M. (2021). Asymmetries in visual acuity around the visual field. Journal of Vision, 21(1), 2.

10. Beaton, A., & Blakemore, C. (1981). Orientation selectivity of the human visual system as a function of retinal eccentricity and visual hemifield. Perception, 10(3), 273–282.

11. Beck, D. M., & Kastner, S. (2014). Neural systems for spatial attention in the human brain: Evidence from neuroimaging in the framework of biased competition. In A. C. Nobre & S. Kastner (Eds.), *Oxford Handbook of Attention*: Oxford University Press.

12. Bell, J., Gheorghiu, E., Hess, R. F., & Kingdom, F. A. (2011). Global shape processing involves a hierarchy of integration stages. Vision Research, 51(15), 1760–1766.

13. Bell, J., Gheorghiu, E., & Kingdom, F. A. A. (2009). Orientation tuning of curvature adaptation reveals both curvature-polarity-selective and non-selective mechanisms. Journal of Vision, 9(12), 3.

14. Benson, N. C., Kupers, E. R., Barbot, A., Carrasco, M., & Winawer, J. (2021). Cortical magnification in human visual cortex parallels task performance around the visual field. Elife, 10, e67685.

15. Binda, P., & Murray, S. O. (2015). Spatial attention increases the pupillary response to light changes. Journal of Vision, 15(2), 1.

16. Boynton, G. M., & Finney, E. M. (2003). Orientation-specific adaptation in human visual cortex. Journal of Neuroscience, 23(25), 8781–8787.

17. Bradley, C., Nydam, A. S., Dux, P. E., & Mattingley, J. B. (2022). State-dependent effects of neural stimulation on brain function and cognition. Nature Reviews Neuroscience, 23(8), 459–475.

18. Brainard, D. H. (1997). The psychophysics toolbox. Spatial Vision, 10(4), 433–436.

19. Buschman, T. J., & Miller, E. K. (2009). Serial, covert shifts of attention during visual search are reflected by the frontal eye fields and correlated with population oscillations. Neuron, 63(3), 386–396.

20. Cameron, E. L., Tai, J. C., & Carrasco, M. (2002). Covert attention affects the psychometric function of contrast sensitivity. Vision Research, 42(8), 949–967.

21. Carrasco, M. (2011). Visual attention: The past 25 years. Vision Research, 51(13), 1484–1525.

22. Carrasco, M. (2014). Spatial covert attention: Perceptual modulation. The Oxford Handbook of Attention, *183*, 230.

23. Carrasco, M., & Barbot, A. (2014). How attention affects spatial resolution. Paper presented at the Cold Spring Harbor Symposia on Quantitative Biology.

24. Carrasco, M., & Barbot, A. (2019). Spatial attention alters visual appearance. Current Opinion in Psychology, 29, 56–64.

25. Carrasco, M., Loula, F., & Ho, Y.-X. (2006). How attention enhances spatial resolution: Evidence from selective adaptation to spatial frequency. Perception & Psychophysics, 68(6), 1004–1012.

26. Carrasco, M., Talgar, C. P., & Cameron, E. L. (2001). Characterizing visual performance fields: Effects of transient covert attention, spatial frequency, eccentricity, task and set size. Spatial Vision, 15(1), 61.

27. Carrasco, M., Williams, P. E., & Yeshurun, Y. (2002). Covert attention increases spatial resolution with or without masks: Support for signal enhancement. Journal of Vision, 2(6), 4.

28. Chakravarthi, R., Papadaki, D., & Krajnik, J. (2022). Visual field asymmetries in numerosity processing. *Attention, Perception*, & Psychophysics, 84(8), 2607–2622.

29. Cheal, M., Lyon, D. R., & Hubbard, D. C. (1991). Does attention have different effects on line orientation and line arrangement discrimination? The Quarterly Journal of Experimental Psychology, 43(4), 825–857.

30. Chen, Q., Lee, H.-H., Hoxha, K., Fernández, A., Hanning, N. M., & Carrasco, M. (2025). Does human right frontal eye field (rFEF+) play a critical role in exogenous attention? A Transcranial Magnetic Stimulation (TMS) study. Journal of Vision, 25(9), 2612–2612.

31. Chica, A. B., Bartolomeo, P., & Lupiáñez, J. (2013). Two cognitive and neural systems for endogenous and exogenous spatial attention. Behavioural Brain Research, 237, 107–123.

32. Corbett, J. E., & Carrasco, M. (2011). Visual performance fields: Frames of reference. PLoS One, *6*(9), e24470.

33. Corbetta, M., Patel, G., & Shulman, G. L. (2008). The reorienting system of the human brain: From environment to theory of mind. Neuron, 58(3), 306–324.

34. Corbetta, M., & Shulman, G. L. (2002). Control of goal-directed and stimulus-driven attention in the brain. Nature Reviews Neuroscience, 3(3), 201–215.

35. Crotty, N., Massa, N., Tellez, D., White, A. L., & Grubb, M. A. (2025). A preliminary investigation of the interaction between expectation and the reflexive allocation of covert spatial attention. Scientific Reports, 15(1), 5778.

36. Curcio, C. A., Sloan Jr, K. R., Packer, O., Hendrickson, A. E., & Kalina, R. E. (1987). Distribution of cones in human and monkey retina: Individual variability and radial asymmetry. Science, 236(4801), 579–582.

37. Curcio, C. A., Sloan, K. R., Kalina, R. E., & Hendrickson, A. E. (1990). Human photoreceptor topography. Journal of Comparative Neurology, 292(4), 497–523.

38. Cutrone, E. K., Heeger, D. J., & Carrasco, M. (2018). On spatial attention and its field size on the repulsion effect. Journal of Vision, 18(6), 8.

39. Dao, D. Y., Lu, Z.-L., & Dosher, B. A. (2006). Adaptation to sine-wave gratings selectively reduces the contrast gain of the adapted stimuli. Journal of Vision, 6(7), 6.

40. Dassanayake, T. L., Michie, P. T., & Fulham, R. (2016). Effect of temporal predictability on exogenous attentional modulation of feedforward processing in the striate cortex. International Journal of Psychophysiology, 105, 9–16.

41. Desimone, R., & Duncan, J. (1995). Neural mechanisms of selective visual attention. Annual Review of Neuroscience, 18(1), 193–222.

42. Dosher, B. A., & Lu, Z.-L. (2000a). Mechanisms of perceptual attention in precuing of location. Vision Research, 40(10), 1269–1292.

43. Dosher, B. A., & Lu, Z.-L. (2000b). Noise exclusion in spatial attention. Psychological Science, 11(2), 139–146.

44. Dugué, L., Merriam, E. P., Heeger, D. J., & Carrasco, M. (2018). Specific visual subregions of TPJ mediate reorienting of spatial attention. Cerebral Cortex, 28(7), 2375–2390.

45. Dugué, L., Merriam, E. P., Heeger, D. J., & Carrasco, M. (2020). Differential impact of endogenous and exogenous attention on activity in human visual cortex. Scientific Reports, 10(1), 21274.

46. Faul, F., Erdfelder, E., Lang, A.-G., & Buchner, A. (2007). G* Power 3: A flexible statistical power analysis program for the social, behavioral, and biomedical sciences. Behavior Research Methods, 39(2), 175–191.

47. Feng, J., & Spence, I. (2017). The effects of spatial endogenous pre-cueing across eccentricities. Frontiers in Psychology, 8, 888.

48. Fernández, A., & Carrasco, M. (2020). Extinguishing exogenous attention via transcranial magnetic stimulation. Current Biology, 30(20), 4078–4084. e4073.

49. Fernández, A., Denison, R. N., & Carrasco, M. (2019). Temporal attention improves perception similarly at foveal and parafoveal locations. Journal of Vision, 19(1), 12.

50. Fernández, A., Hanning, N. M., & Carrasco, M. (2023). Transcranial magnetic stimulation to frontal but not occipital cortex disrupts endogenous attention. Proceedings of the National Academy of Sciences, 120(10), e2219635120.

51. Fernández, A., Okun, S., & Carrasco, M. (2022). Differential effects of endogenous and exogenous attention on sensory tuning. Journal of Neuroscience, 42(7), 1316–1327.

52. Fiebelkorn, I. C., & Kastner, S. (2020). Functional specialization in the attention network. Annual Review of Psychology, 71(1), 221–249.

53. Fuller, S., & Carrasco, M. (2009). Perceptual consequences of visual performance fields: The case of the line motion illusion. Journal of Vision, 9(4), 13.

54. Fuller, S., Rodriguez, R. Z., & Carrasco, M. (2008). Apparent contrast differs across the vertical meridian: Visual and attentional factors. Journal of Vision, 8(1), 16.

55. Gao, Y., Webster, M. A., & Jiang, F. (2019). Dynamics of contrast adaptation in central and peripheral vision. Journal of Vision, 19(6), 23.

56. Gardner, J. L., Sun, P., Waggoner, R. A., Ueno, K., Tanaka, K., & Cheng, K. (2005). Contrast adaptation and representation in human early visual cortex. Neuron, 47(4), 607–620.

57. Geweke, F., Pokta, E., & Störmer, V. S. (2021). Spatial distance of target locations affects the time course of both endogenous and exogenous attentional deployment. Journal of Experimental Psychology: Human Perception and Performance, 47(6), 774.

58. Giordano, A. M., McElree, B., & Carrasco, M. (2009). On the automaticity and flexibility of covert attention: A speed-accuracy trade-off analysis. Journal of Vision, 9(3), 30.

59. Greenlee, M. W., Georgeson, M. A., Magnussen, S., & Harris, J. P. (1991). The time course of adaptation to spatial contrast. Vision Research, 31(2), 223–236.

60. Greenwood, J. A., Szinte, M., Sayim, B., & Cavanagh, P. (2017). Variations in crowding, saccadic precision, and spatial localization reveal the shared topology of spatial vision. Proceedings of the National Academy of Sciences, 114(17), E3573–E3582.

61. Grubb, M. A., Behrmann, M., Egan, R., Minshew, N. J., Carrasco, M., & Heeger, D. J. (2013). Endogenous spatial attention: Evidence for intact functioning in adults with autism. Autism Research, 6(2), 108–118.

62. Hanning, N. M., Himmelberg, M. M., & Carrasco, M. (2022). Presaccadic attention enhances contrast sensitivity, but not at the upper vertical meridian. Iscience, 25(2).

63. Hanning, N. M., Himmelberg, M. M., & Carrasco, M. (2024). Presaccadic attention depends on eye movement direction and is related to V1 cortical magnification. Journal of Neuroscience, 44(12).

64. Hein, E., Rolke, B., & Ulrich, R. (2006). Visual attention and temporal discrimination: Differential effects of automatic and voluntary cueing. Visual Cognition, 13(1), 29–50.

65. Herrmann, K., Montaser-Kouhsari, L., Carrasco, M., & Heeger, D. J. (2010). When size matters: Attention affects performance by contrast or response gain. Nature Neuroscience, 13(12), 1554–1559.

66. Himmelberg, M. M., Kurzawski, J. W., Benson, N. C., Pelli, D. G., Carrasco, M., & Winawer, J. (2021). Cross-dataset reproducibility of human retinotopic maps. Neuroimage, 244, 118609.

67. Himmelberg, M. M., Kwak, Y., Carrasco, M., & Winawer, J. (2025). Unpacking the V1 map: Differential covariation of preferred spatial frequency and cortical magnification across spatial dimensions. PLOS Computational Biology, 21(10), e1013599.

68. Himmelberg, M. M., Tünçok, E., Gomez, J., Grill-Spector, K., Carrasco, M., & Winawer, J. (2023). Comparing retinotopic maps of children and adults reveals a late-stage change in how V1 samples the visual field. Nature Communications, 14(1), 1561.

69. Himmelberg, M. M., Winawer, J., & Carrasco, M. (2020). Stimulus-dependent contrast sensitivity asymmetries around the visual field. Journal of Vision, 20(9), 18.

70. Himmelberg, M. M., Winawer, J., & Carrasco, M. (2022). Linking individual differences in human primary visual cortex to contrast sensitivity around the visual field. Nature Communications, 13(1), 3309.

71. Himmelberg, M. M., Winawer, J., & Carrasco, M. (2023). Polar angle asymmetries in visual perception and neural architecture. Trends in Neurosciences, 46(6), 445–458.

72. Hopfinger, J. B., Luck, S. J., & Hillyard, S. A. (2004). Selective attention: Electrophysiological and neuromagnetic studies. The Cognitive Neurosciences, 3, 561–574.

73. Hopfinger, J. B., & West, V. M. (2006). Interactions between endogenous and exogenous attention on cortical visual processing. NeuroImage, 31(2), 774–789.

74. Huang, D., Liao, K., Chen, F., & Chen, Y. (2025). The effects of acute alcohol intake on endogenous vs. exogenous attention in visual perception. Addictive Behaviors, 108465.

75. Jigo, M., & Carrasco, M. (2018). Attention alters spatial resolution by modulating second-order processing. Journal of Vision, 18(7), 2.

76. Jigo, M., & Carrasco, M. (2020). Differential impact of exogenous and endogenous attention on the contrast sensitivity function across eccentricity. Journal of Vision, 20(6), 11.

77. Jigo, M., Heeger, D. J., & Carrasco, M. (2021). An image-computable model of how endogenous and exogenous attention differentially alter visual perception. Proceedings of the National Academy of Sciences, 118(33), e2106436118.

78. Jigo, M., Tavdy, D., Himmelberg, M. M., & Carrasco, M. (2023). Cortical magnification eliminates differences in contrast sensitivity across but not around the visual field. eLife, 12, e84205.

79. Kastner, S., & Buschman, T. J. (2017). Visual attention. In *Oxford Research Encyclopedia of Neuroscience*.

80. Kim, C., & Chong, S. C. (2024). Metacognition of perceptual resolution across and around the visual field. Cognition, 253, 105938.

81. Kinchla, R. (1980). The measurement of attention. In R. S. Nikerson (Ed.), Attention and performance VIII (pp. 213–238): Psychology Press.

82. Kobayashi, M., & Pascual-Leone, A. (2003). Transcranial magnetic stimulation in neurology. The Lancet Neurology, 2(3), 145–156.

83. Kohn, A. (2007). Visual adaptation: Physiology, mechanisms, and functional benefits. Journal of Neurophysiology, 97(5), 3155–3164.

84. Kovács, G., Zimmer, M., Harza, I., & Vidnyánszky, Z. (2007). Adaptation duration affects the spatial selectivity of facial aftereffects. Vision Research, 47(25), 3141–3149.

85. Kupers, E. R., Benson, N. C., Carrasco, M., & Winawer, J. (2022). Asymmetries around the visual field: From retina to cortex to behavior. PLoS Computational Biology, 18(1), e1009771.

86. Kurzawski, J. W., Burchell, A., Thapa, D., Winawer, J., Majaj, N. J., & Pelli, D. G. (2023). The Bouma law accounts for crowding in 50 observers. Journal of Vision, 23(8), 6.

87. Kwak, Y., Hanning, N. M., & Carrasco, M. (2023). Presaccadic attention sharpens visual acuity. Scientific Reports, 13(1), 2981.

88. Kwak, Y., Hanning, N. M., & Carrasco, M. (in press). Saccade direction modulates the temporal dynamics of presaccadic attention. Journal of Vision.

89. Kwak, Y., Zhao, Y., Lu, Z.-L., Hanning, N. M., & Carrasco, M. (2024). Presaccadic attention enhances and reshapes the Contrast Sensitivity Function differentially around the visual field. eNeuro, 11(9).

90. Larsson, J., & Harrison, S. J. (2015). Spatial specificity and inheritance of adaptation in human visual cortex. Journal of Neurophysiology, 114(2), 1211–1226.

91. Lauritzen, T. Z., D’Esposito, M., Heeger, D. J., & Silver, M. A. (2009). Top–down flow of visual spatial attention signals from parietal to occipital cortex. Journal of Vision, 9(13), 18.

92. Lee, H.-H., & Carrasco, M. (2025). Visual adaptation stronger at the horizontal than the vertical meridian: Linking performance with V1 cortical surface area. Proceedings of the National Academy of Sciences, 122(29), e2507810122.

93. Lee, H.-H., Fernández, A., & Carrasco, M. (2024). Adaptation and exogenous attention interact in the early visual cortex: A TMS study. Iscience, 27(11).

94. Lennie, P. (2003). The cost of cortical computation. Current Biology, 13(6), 493–497.

95. Li, H.-H., Pan, J., & Carrasco, M. (2021). Different computations underlie overt presaccadic and covert spatial attention. Nature Human Behaviour, 5(10), 1418–1431.

96. Lin, Y., Zhou, X., Naya, Y., Gardner, J. L., & Sun, P. (2021). Voxel-wise linearity analysis of increments and decrements in BOLD responses in human visual cortex using a contrast adaptation paradigm. Frontiers in Human Neuroscience, 15, 541314.

97. Ling, S., & Carrasco, M. (2006a). Sustained and transient covert attention enhance the signal via different contrast response functions. Vision Research, 46(8-9), 1210–1220.

98. Ling, S., & Carrasco, M. (2006b). When sustained attention impairs perception. Nature Neuroscience, 9(10), 1243–1245.

99. Ling, S., Jehee, J. F., & Pestilli, F. (2015). A review of the mechanisms by which attentional feedback shapes visual selectivity. Brain Structure and Function, 220(3), 1237–1250.

100. Liu, T., Pestilli, F., & Carrasco, M. (2005). Transient attention enhances perceptual performance and fMRI response in human visual cortex. Neuron, 45(3), 469–477.

101. Liu, T., Stevens, S. T., & Carrasco, M. (2007). Comparing the time course and efficacy of spatial and feature-based attention. Vision Research, 47(1), 108–113.

102. Lu, Z.-L., Lesmes, L. A., & Dosher, B. A. (2002). Spatial attention excludes external noise at the target location. Journal of Vision, 2(4), 4.

103. Luck, S. J., & Thomas, S. J. (1999). What variety of attention is automatically captured by peripheral cues? Perception & Psychophysics, 61(7), 1424–1435.

104. Luzardo, F., & Yeshurun, Y. (2025). Large-scale examination of the benefit and cost of spatial attention and their individual variability. Cognition, 264, 106242.

105. Mangun, G., & Hillyard, S. (1990). Allocation of visual attention to spatial locations: Tradeoff functions for event-related brain potentials and detection performance. Perception & Psychophysics, 47(6), 532–550.

106. Meyer, K. N., Du, F., Parks, E., & Hopfinger, J. B. (2018). Exogenous vs. endogenous attention: Shifting the balance of fronto-parietal activity. Neuropsychologia, 111, 307–316.

107. Montaser-Kouhsari, L., & Carrasco, M. (2009). Perceptual asymmetries are preserved in short-term memory tasks. *Attention, Perception*, & Psychophysics, 71(8), 1782–1792.

108. Montaser-Kouhsari, L., & Rajimehr, R. (2004). Attentional modulation of adaptation to illusory lines. Journal of Vision, 4(6), 3.

109. Morrone, M. C., Denti, V., & Spinelli, D. (2004). Different attentional resources modulate the gain mechanisms for color and luminance contrast. Vision Research, 44(12), 1389–1401.

110. Nakayama, K., & Mackeben, M. (1989). Sustained and transient components of focal visual attention. Vision Research, 29(11), 1631–1647.

111. Olivers, C. N. (2025). Selective attention and eccentricity: A comprehensive review. Neuroscience & Biobehavioral Reviews, 106368.

112. Palmieri, H., & Carrasco, M. (2024). Task demand mediates the interaction of spatial and temporal attention. Scientific Reports, 14(1), 9228.

113. Pelli, D. G. (1997). The VideoToolbox software for visual psychophysics: transforming numbers into movies. Spatial Vision, 10(4), 437–442.

114. Perini, F., Cattaneo, L., Carrasco, M., & Schwarzbach, J. V. (2012). Occipital transcranial magnetic stimulation has an activity-dependent suppressive effect. Journal of Neuroscience, 32(36), 12361–12365.

115. Pestilli, F., & Carrasco, M. (2005). Attention enhances contrast sensitivity at cued and impairs it at uncued locations. Vision Research, 45(14), 1867–1875.

116. Pestilli, F., Carrasco, M., Heeger, D. J., & Gardner, J. L. (2011). Attentional enhancement via selection and pooling of early sensory responses in human visual cortex. Neuron, 72(5), 832–846.

117. Pestilli, F., Ling, S., & Carrasco, M. (2009). A population-coding model of attention’s influence on contrast response: Estimating neural effects from psychophysical data. Vision Research, 49(10), 1144–1153.

118. Pestilli, F., Viera, G., & Carrasco, M. (2007). How do attention and adaptation affect contrast sensitivity? Journal of Vision, 7(7), 9.

119. Petrov, Y., & Meleshkevich, O. (2011). Asymmetries and idiosyncratic hot spots in crowding. Vision Research, 51(10), 1117–1123.

120. Prins, N., & Kingdom, F. A. (2018). Applying the model-comparison approach to test specific research hypotheses in psychophysical research using the Palamedes toolbox. Frontiers in Psychology, 9, 1250.

121. Purokayastha, S., Roberts, M., & Carrasco, M. (2021). Voluntary attention improves performance similarly around the visual field. *Attention, Perception*, & Psychophysics, 83(7), 2784–2794.

122. Ramamurthy, M., White, A. L., & Yeatman, J. D. (2024). Children with dyslexia show no deficit in exogenous spatial attention but show differences in visual encoding. Developmental Science, 27(3), e13458.

123. Remington, R. W., Johnston, J. C., & Yantis, S. (1992). Involuntary attentional capture by abrupt onsets. Perception & Psychophysics, 51(3), 279–290.

124. Reynolds, J. H., & Heeger, D. J. (2009). The normalization model of attention. Neuron, 61(2), 168–185.

125. Roberts, M., Ashinoff, B. K., Castellanos, F. X., & Carrasco, M. (2018). When attention is intact in adults with ADHD. Psychonomic Bulletin & Review, 25(4), 1423–1434.

126. Roberts, M., Cymerman, R., Smith, R. T., Kiorpes, L., & Carrasco, M. (2016). Covert spatial attention is functionally intact in amblyopic human adults. Journal of Vision, 16(15), 30.

127. Schieting, S., & Spillmann, L. (1987). Flicker adaptation in the peripheral retina. Vision Research, 27(2), 277–284.

128. Silva, M. F., Brascamp, J. W., Ferreira, S., Castelo-Branco, M., Dumoulin, S. O., & Harvey, B. M. (2018). Radial asymmetries in population receptive field size and cortical magnification factor in early visual cortex. NeuroImage, 167, 41–52.

129. Song, H., Chui, T. Y. P., Zhong, Z., Elsner, A. E., & Burns, S. A. (2011). Variation of cone photoreceptor packing density with retinal eccentricity and age. Investigative Ophthalmology & Visual Science, 52(10), 7376–7384.

130. Sperling, G., & Melchner, M. J. (1978). The attention operating characteristic: Examples from visual search. Science, 202(4365), 315–318.

131. Talgar, C. P., & Carrasco, M. (2002). Vertical meridian asymmetry in spatial resolution: Visual and attentional factors. Psychonomic Bulletin & Review, 9(4), 714–722.

132. Team, R. C. (2000). R language definition. *Vienna, Austria: R foundation for statistical computing*, *3*(1), 116.

133. Tsai, L.-T., Liao, K.-M., Hou, C.-H., Jang, Y., & Chen, C.-C. (2024). Visual field asymmetries in visual word form identification. Vision Research, 220, 108413.

134. Tünçok, E., Carrasco, M., & Winawer, J. (2025). How spatial attention alters visual cortical representation during target anticipation. Nature Communications, 16, 8746.

135. Tünçok, E., Kiorpes, L., & Carrasco, M. (2025). Opposite asymmetry in visual perception of humans and macaques. Current Biology, 35(3), 681–687. e684.

136. Valero-Cabré, A., Pascual-Leone, A., & Coubard, O. A. (2011). Transcranial magnetic stimulation (TMS) in basic and clinical neuroscience research. Revue Neurologique, 167(4), 291–316.

137. Vergeer, M., Mesik, J., Baek, Y., Wilmerding, K., & Engel, S. A. (2018). Orientation-selective contrast adaptation measured with SSVEP. Journal of Vision, 18(5), 2.

138. Vinke, L. N., Bloem, I. M., & Ling, S. (2022). Saturating nonlinearities of contrast response in human visual cortex. Journal of Neuroscience, 42(7), 1292–1302.

139. Wang, F., Chen, M., Yan, Y., Zhaoping, L., & Li, W. (2015). Modulation of neuronal responses by exogenous attention in macaque primary visual cortex. Journal of Neuroscience, 35(39), 13419–13429.

140. Wang, Z., Murai, Y., & Whitney, D. (2020). Idiosyncratic perception: A link between acuity, perceived position and apparent size. Proceedings of the Royal Society B, 287(1930), 20200825.

141. Webster, M. A. (2011). Adaptation and visual coding. Journal of Vision, 11(5), 3.

142. Webster, M. A. (2015). Visual adaptation. Annual Review of Vision Science, 1(1), 547–567.

143. Westerberg, J. A., Schall, J. D., Woodman, G. F., & Maier, A. (2023). Feedforward attentional selection in sensory cortex. Nature Communications, 14(1), 5993.

144. Xue, S., Barbot, A., Abrams, J., Chen, Q., & Carrasco, M. (2025). System-Level computations underlie visual field heterogeneity. bioRxiv. 10.1101/2025.09.19.677418

145. Yantis, S., & Jonides, J. (1996). Attentional capture by abrupt onsets: new perceptual objects or visual masking? Journal of Experimental Psychology: Human Perception and Performance, 22(6), 1505–1513.

146. Yeshurun, Y., & Carrasco, M. (1998). Attention improves or impairs visual performance by enhancing spatial resolution. Nature, 396(6706), 72–75.

147. Yeshurun, Y., Montagna, B., & Carrasco, M. (2008). On the flexibility of sustained attention and its effects on a texture segmentation task. Vision Research, 48(1), 80–95.

148. Zimmermann, E., Weidner, R., Abdollahi, R. O., & Fink, G. R. (2016). Spatiotopic adaptation in visual areas. Journal of Neuroscience, 36(37), 9526–9534.

